# Decreased social, drug, and food reward sensitivity in adolescent mice

**DOI:** 10.1101/2025.08.18.670792

**Authors:** Klaudia Misiołek, Magdalena Chrószcz, Marta Klimczak, Aleksandra Rzeszut, Julia Netczuk, Barbara Ziółkowska, Łukasz Szumiec, Maria Kaczmarczyk-Jarosz, Zofia Harda, Jan Rodriguez Parkitna

**Affiliations:** Department of Molecular Neuropharmacology, Maj Institute of Pharmacology, Polish Academy of Sciences, Krakow, Poland; Department of Physiology, Maj Institute of Pharmacology, Polish Academy of Sciences, Krakow, Poland

## Abstract

Adolescence shapes adaptive adult behaviors. Most studies compare single adolescent and adult age groups using only one type of reward, limiting insight into developmental trajectories underlying behavioral change. Here, we investigated how social, cocaine, and palatable food rewards become associated with environmental contexts in female and male C57BL/6 mice across early- (pubertal onset), mid- (peripubertal phase), and late- (sexual maturity) adolescence, compared to adults. Using the conditioned place preference (CPP) task, we found that generally all rewards induced place preference for the reward-associated context, with only minor effects of sex. In contrast, age significantly influenced CPP expression. Adolescent mice exhibited a significantly reduced CPP compared to adults in palatable food and social CPP paradigms, evident in both decreased mean conditioned context preference and lower proportion of animals that developed a preference after conditioning. Cocaine CPP was not significantly affected by age. Direct comparisons across CPP task outcomes further confirmed that age, rather than reward type or sex, was the primary factor influencing the magnitude of CPP. Specifically, mid- and late-adolescent mice showed reduced mean reward CPP, and mid-adolescents were less likely to express a reward preference relative to adults. Based on the behavioral analyses, we conclude that the lower expression of preference for a conditioned context in adolescent animals is due to developmental changes in reward sensitivity, rather than deficits in learning or higher novelty-seeking behavior.

## Introduction

Adolescence is a crucial transitional phase leading to adulthood, characterized by significant physical, hormonal, and behavioral development. It features increased sensitivity to environmental factors and heightened neural plasticity in brain regions responsible for learning, motivation, emotional regulation, and social cognition [1–5]. While this extensive maturation promotes adaptive growth, it also creates vulnerability to maladaptive changes, particularly under stress, which may trigger early onset of psychiatric disorders [4,6]. A key factor in this process is the development of the brain’s reward system, connected to increased reward-seeking behavior [3,7,8], novelty or sensation seeking [4,9], risk-taking [10,11], and susceptibility to drugs of abuse [12,13]. It has been consistently reported that gender significantly influences reward-related behaviors [14–17] as well as the onset, prevalence, and symptoms of mental disorders [18]. Teenage girls and boys are more sensitive to social context, process social rewards differently than adults, and generally show heightened sensitivity to all rewards within peer settings [4,19,20]. The subjective valuation of rewards undergoes significant changes across development [21–23]; however, it remains uncertain whether sensitivity to various rewarding stimuli (such as social, food, or drug) follows a common developmental pattern. Heightened neural responsiveness to socially salient stimuli in adolescents is accompanied by increased risk-taking behaviors in social contexts, such as initial substance use, potentially elevating the risk for mental disorders [4,19,20]. This relationship suggests that social reward sensitivity may develop differently from non-social reward processing.

Adolescence and its associated behaviors show notable similarities between humans and laboratory rodents. The ages from 10 to 19 years in humans correspond to postnatal days 28 to 55 in mice, with both species experiencing similar developmental milestones, including the maturation of the brain’s reward system [10,24]. It has been reported that adolescent mice may have increased sensitivity to the rewarding effects of drugs of abuse [25], especially under stress conditions [26,27], and are less sensitive to their aversive effects [13]. Adolescent, but not adult mice, were observed to readily develop a preference for a social-conditioned context [28] and demonstrate a stronger motivation to obtain highly palatable food [29–32]. These findings consistently support the idea that increased reward sensitivity during adolescence may strengthen associative memory, enhance the salience of related cues, and simultaneously facilitate the acquisition of both adaptive and maladaptive behaviors driven by environmental factors such as social experiences, drugs of abuse, or diet. Although this hypothesis is widely accepted, direct empirical evidence remains limited. In both humans and rodents, developmental processes are uneven and dynamic; females and males mature at different rates. Consequently, incorporating these factors into experiments is essential to capture and understand behavioral changes accurately. Nardou and colleagues demonstrated that conditioned place preference for social reward peaks during late adolescence compared to adulthood in both male and female mice [28]. Additionally, a study on rats reported a sex-independent decline in social reward motivation during mid-adolescence, with both studies emphasizing the importance of multi-timepoint assessments throughout adolescence [33]. Equivalent analyses for other types of rewards are currently lacking. Moreover, despite the common view that adolescence is characterized by heightened reward sensitivity, it was historically debated whether a reduction in reward sensitivity could actually drive increased reward-seeking behaviors, potentially explaining vulnerability to drug abuse in humans [4].

The conditioned place preference (CPP) paradigm is a fundamental method for assessing the rewarding properties of a stimulus. Originally developed to evaluate the reinforcing effects of drugs [34–36], it has been adapted to examine various stimuli, including pain, food and social cues. In this study, we used modified CPP tasks to measure reward-conditioned preference for social contact, palatable food, or cocaine in male and female C57BL/6 mice at three stages of adolescence: early (∼post-natal days 33, indicating puberty onset), mid (∼P38, peripubertal period), and late (∼P43, reflecting sexual maturity). Unexpectedly, we found that the expression of reward-conditioned preference is typically lower in mid-adolescents compared to adult animals, regardless of reward type or sex. This finding challenge the simplistic notion that heightened reward sensitivity during adolescence directly leads to a strengthening of reward-context associations.

## Materials and Methods

### Animals

Experiments were conducted on male and female C57BL/6 mice bred at the animal facility of the Maj Institute of Pharmacology, Polish Academy of Sciences in Krakow. The animals were housed at 22 ± 2°C, with 40-60% humidity, and maintained on a 12/12 hour light/dark cycle (lights on at 7 AM). Mice had unlimited access to water and food (maintenance chow, 10mm pellets, Altromin Spezialfutter, cat no. 1324, Germany). After weaning, they were housed in groups of 2-6 littermates per cage in standard Plexiglas cages (length 325 mm × width 170 mm) with wooden blocks for gnawing and nesting material. Female and male mice were kept in separate rooms. All behavioral procedures were performed during the light phase under dim light conditions (5-10 lux) with infrared lighting. Mice were handled for 3-5 consecutive days before each experiment by placing them in the experimenter’s hands for 2-3 minutes. Every 3 days, animals were marked on the tail for identification. Animals were moved to the experimental room at least 30 minutes before the experiments began. The age, sex, and weight of each animal at the start of the experiments are listed in **Table S1**. All procedures were approved by the 2nd Local Bioethics Committee in Krakow (permit numbers 293/2020, 32/2021, 55/2024, 231/2024) and adhered to the guidelines of the European Parliament and the Council of 22 September 2010, on the protection of animals used for scientific purposes (Directive 2010/63/EU and Polish Law Dz.U. 2015 poz. 266). The experiments were planned and reported in accordance with ARRIVE guidelines [37].

### Social conditioned place preference

Social CPP was performed as described before [38–41] and is a revised version of the protocol by Panksepp and Lahvis [42], with modifications introduced by Dölen and colleagues [28,43]. Compared to the procedure we described previously, the main change is the bedding types. Animals were housed on corn bedding prior to experiments (corn 2 mm, Rehofix MK2000, Germany). The test was conducted in a custom-made two-chamber apparatus with compartments differing in bedding type (context A: aspen, ABEDD, Latvia or Tapvei GLP, Estonia, context B: 1/8’ Pelleted Cellulose, Scott Pharma Solutions, cat no. L0107, USA) and wooden blocks (context A: cuboid, context B: cube, both from Zoolab, Poland). The social CPP involved three phases: pre-test, conditioning, and post-test (**Fig. 1A**). Behavior during pre- and post-test was recorded with a camera (acA1300 – 60 gm, Basler, Germany). During pre-test, mice freely explored the two-chamber cages for 30 min. Mice that spent more than 70% of the time in any context were excluded. To counterbalance contextual bias, half of the animals were assigned to context A as the social-paired environment, and the other half to context B. The conditioning phase included 6 sessions on consecutive days. During the social conditioning session, mice and their littermates were moved to new home cages with one of the conditioning contexts for 24 hours. During the isolation session, mice were placed separately in new home cages with the other conditioning context for 24 hours. Conditioning started with a social session, and the context alternated daily (social-isolation-social-isolation-social-isolation). The post-test was conducted the day after the last conditioning session. Behavior was automatically analyzed using EthoVision XT 15 software (Noldus, The Netherlands). The social place preference index was calculated by subtracting the time spent in the social context during the pre-test from the time spent in the social context during the post-test.

**Fig 1.**
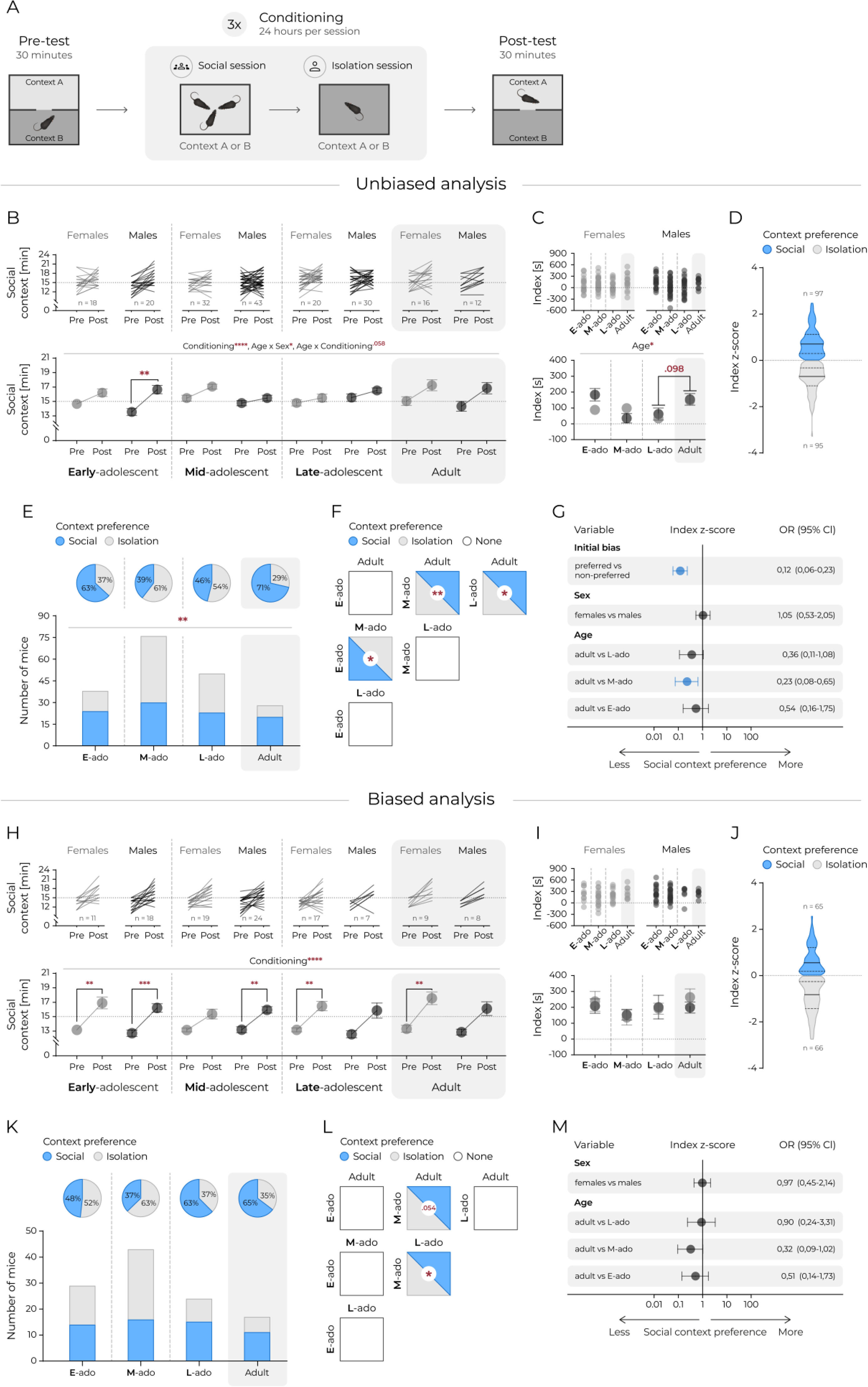
Preference for the context associated with social cues is influenced by age and initial context bias, but not by sex. **(A)** A schematic representation of the social CPP. **(B)** Time spent in the social context. Top panels: lines represent individual animals. Bottom panel: mean values. Circles and connecting lines represent means and matched values. Whiskers represent s.e.m. values. Dotted lines represent random value (i.e. 15 min). Females and males are shown in gray and black respectively. Statistical analysis was performed using 3-way ANOVA and post hoc Tukey’s HSD, ‘**’ corresponds to p ≤ .01, ‘***’, p ≤ .001, ‘****’ p ≤ .0001. **(C)** Difference in seconds between time spent in social context posttest and time spent in social context pretest (index). Top panels: individual animals. Bottom panel: mean values. Whiskers represent s.e.m. values. Dotted lines represent no change. Statistical analysis was performed using 2-way ANOVA and post hoc Tukey’s HSD, *’ corresponds to p ≤ .05. **(D)** Distribution of context preference (social or isolation) in posttest. Sex and age data were combined. Index z-score calculated as: (index of individual animal – mean index of all mice) / standard deviation of index of all mice). Context preference interpreted according to index z-score value: positive, i.e. > 0 – “social” or negative i.e. < 0 – “isolation”. Solid line represents the median, dashed lines mark the quartiles. **(E)** Fractions of animals showing increase in preference of social context. Top panels (pie charts): percent of animals expressing social (blue) or isolation (white) context preference in each age group. Bottom panels: proportion of animals expressing social (blue bars) or isolation (white bars) context preference. χ^2^ test revealed no significant differences between proportions. **(F)** Pairwise comparisons of the fraction of animals with increased social context preference at different development stages. Each square represents a comparison between each age group. The color of the square fill indicates the predominance of the proportion of animals expressing social (blue), isolation (gray) or none (white) preferences of one group over another. All pairwise χ^2^ comparisons were performed, ‘*’ corresponds to p ≤ .05, ‘**’ corresponds to p ≤ .01. **(G)** Estimated effects of initial bias, sex and age on the odds of increased preference for social context based on logistic regression. The circles and horizontal lines indicate the odds ratio and corresponding 95% confidence intervals. Statistically significant effects are marked blue. Panels H – M correspond to B – G but with analyses performed only on animals initially preferring the isolation context, i.e. the equivalent of a biased experimental design. **(H)** Time spent in the social context. Top panels: lines represent individual animals. Bottom panel: mean values. Whiskers represent s.e.m. values. Dotted lines represent random value (i.e. 15 min). Females and males are shown in gray and black respectively. Statistical analysis was performed using 3-way ANOVA and post hoc Tukey’s HSD, **’ corresponds to p ≤ .01, ‘***’ p ≤ .001, ‘****’ p ≤ .0001. **(I)** Difference in seconds between time spent in social context posttest and time spent in social context pretest (index). Top panels: individual animals. Bottom panel: mean values. Circles and connecting lines represent means and matched values. Whiskers represent s.e.m. values. Dotted lines represent no change. Statistical analysis was performed using 2-way ANOVA revealed no significant differences between groups. **(J)** Distribution of context preference (social or isolation) in posttest. Sex and age data were combined. Index z-score calculated as: (index of individual animal – mean index of all mice) / standard deviation of index of all mice). Context preference interpreted according to index z-score value: positive, i.e. > 0 – “social” or negative, i.e. < 0 – “isolation”. Solid line represents the median, dashed lines mark the quartiles. **(K)** Fractions of animals showing increase in preference of social context. Top panels (pie charts): percent of animals expressing social (blue) or isolation (white) context preference in each age group. Bottom panels: proportion of animals expressing social (blue bars) or isolation (white bars) context preference. χ^2^ test revealed no significant differences between proportions. **(L)** Pairwise comparisons of the fraction of animals with increased social context preference at different development stages. Each square represents a comparison between each age group. The color of the square fill indicates the predominance of the proportion of animals expressing social (blue), isolation (gray) or none (white) preferences of one group over another. All pairwise χ^2^ comparisons were performed, ‘*’ corresponds to p ≤ .05. **(M)** Estimated effects of sex and age on the odds of increased preference for social context based on logistic regression. The circles and horizontal lines indicate the odds ratio and corresponding 95% confidence intervals. There were no statistically significant effects, with one result approaching statistical significance adult vs. M-ado, p = .059.

### Cocaine conditioned place preference

The cocaine CPP test was conducted using an automatic three-chamber apparatus (Med Associates, St. Albans, VT, USA, MED-CPP-MSAT) as described previously [39,44,45]. The apparatus features two compartments that differ in color and tactile cues, along with a middle compartment with guillotine doors between them. Photobeams automatically tracked movement and time spent in each chamber. The CPP paradigm included three phases: pre-test, conditioning, and post-test (**Fig. 2A**). On the first day, the pre-test was performed to determine initial preference. Mice were placed in the middle compartment and allowed to explore the entire apparatus for 20 minutes. Mice that spent more than 70% of their time in one of the conditioning chambers, excluding time in the middle chamber, were excluded. Conditioning sessions took place over the next three days. Each conditioning day included two 40-minute sessions: one with saline, and after about 4 hours in home cages, a cocaine session (saline-cocaine, saline-cocaine, saline-cocaine). The design was biased; the less preferred chamber was paired with the cocaine injection (cocaine hydrochloride dissolved in saline, i.p., 10 mg/kg, 5 μl/g, cocaine hydrochloride, Toronto Research Chemicals; TRC, Toronto, North York, ON, Canada), while the more preferred chamber was paired with the saline injection (i.p., 5 μl/g, Polpharma). The post-test was performed in the same way as the pre-test the day after the last conditioning session. The place preference index was calculated by subtracting the time spent in the cocaine-paired chamber during the pre-test from the time spent in the cocaine-paired chamber during the post-test.

**Fig 2.**
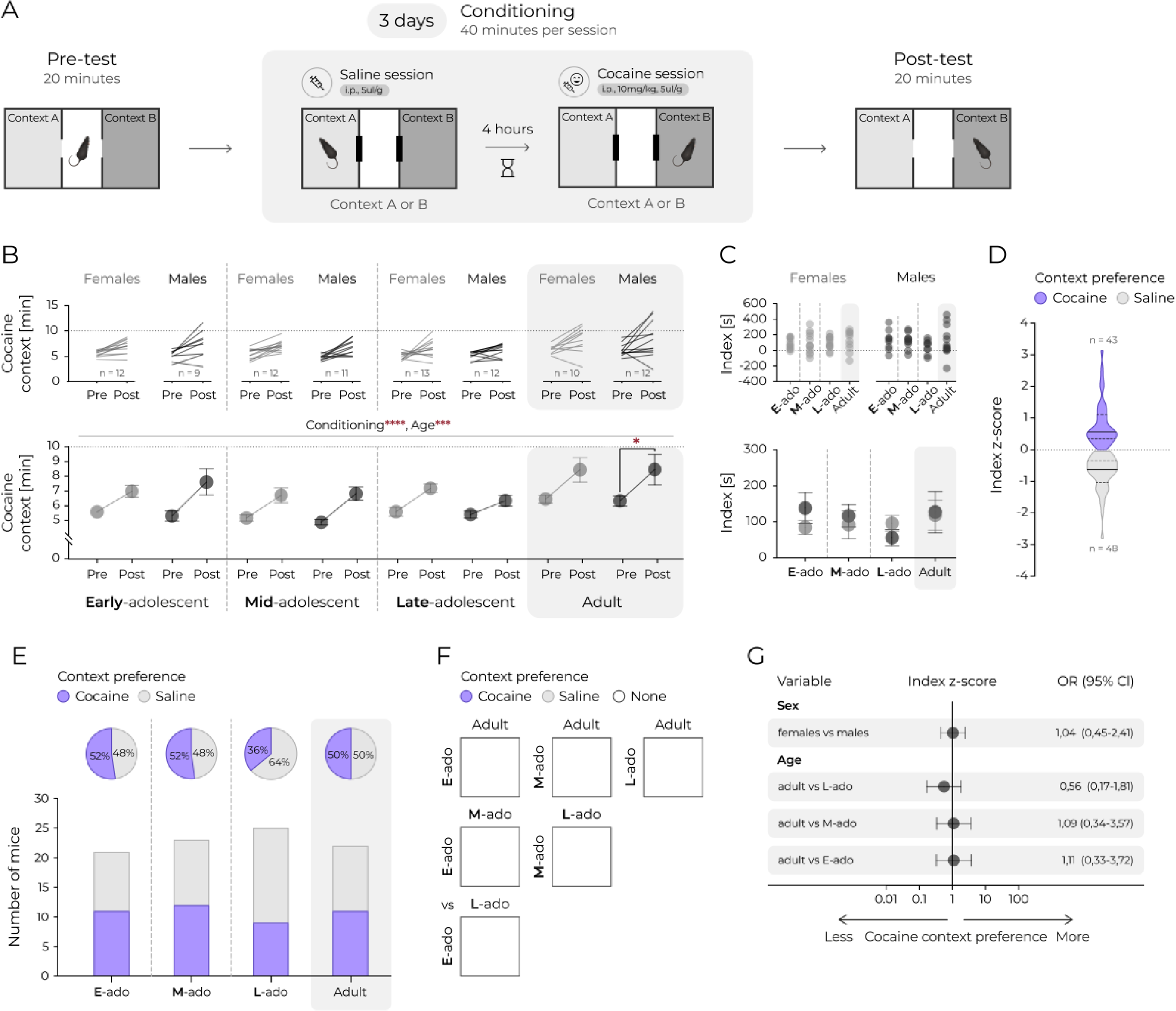
Adolescent and adult mice show similar preferences for the context associated with cocaine administration. **(A)** A schematic representation of the cocaine CPP. **(B)** Time spent in the cocaine context. Top panels: lines represent individual animals. Bottom panel: mean values. Circles and connecting lines represent means and matched values. Whiskers represent s.e.m. values. Dotted lines represent random value (i.e. 10 min). Females and males are shown in gray and black, respectively. Statistical analysis was performed using 3-way ANOVA and post hoc Tukey’s HSD, ‘***’ corresponds to p ≤ .001, ‘****’ p ≤ .0001. **(C)** Difference in seconds between time spent in cocaine context posttest and time spent in saline context pretest (index). Top panels: individual animals. Bottom panel: mean values. Whiskers represent s.e.m. values. Dotted lines represent no change. Statistical analysis was performed using 2-way ANOVA and revealed no significant differences between groups. **(D)** Distribution of context preference (cocaine or saline) in posttest. Sex and age data were combined. Index z-score calculated as: (index of individual animal – mean index of all mice) / standard deviation of index of all mice. Context preference interpreted according to index z-score value: positive– “cocaine” or negative, “saline”. The solid line represents the median, and the dashed lines mark the quartiles. **(E)** Fractions of animals showing an increase in preference for the cocaine context. Top panels (pie charts): percent of animals expressing cocaine (purple) or saline (white) context preference in each age group. Bottom panels: proportion of animals expressing cocaine (purple bars) or saline (white bars) context preference. χ^2^ test revealed no significant differences between proportions. **(F)** Pairwise comparisons of the fraction of animals with increased cocaine context preference at different development stages. Each square represents a comparison between each age group. The color of the square fill indicates the predominance of the proportion of animals expressing cocaine (purple), saline (gray) or none (white) preferences of one group over another. All pairwise χ^2^ comparisons were not significant. **(G)** Estimated effects of sex and age on the odds of increased preference for social context based on logistic regression. The circles and horizontal lines indicate the odds ratio and corresponding 95% confidence intervals. There were no statistically significant effects.

### Palatable food conditioned place preference

A palatable food CPP test was conducted in the same experimental setup as the cocaine CPP, using a protocol adapted from Clough and colleagues [46]. Mice were housed individually and habituated to a palatable food mixture placed in their home cage on a flat glass plate. Each day, they received a fresh portion consisting of 2 pieces of regular chow, 1/3 of an open Oreos cookie (cream side up), 2 Cheetos, and 5 Froot Loops; the nutritional details of these foods are listed in **Table S2**. This habituation occurred for two days before the experiment. On the pre-test day, mice were grouped by sex with their littermates in their original home cages (without palatable food, but with chow and water available freely) and allowed to habituate in the experimental room. The pre-test was identical to the cocaine CPP test procedure (**Fig. 3A**). Mice that spent over 70% of their time (excluding time in the middle box) in any conditioning box were excluded from the analysis. An initial bias test was performed, and a biased design was used to assign the less preferred context as the palatable food context (with the food mix) and the more preferred one as the empty cage (without food). Conditioning sessions lasted six consecutive days, with one 60-minute session each day. To motivate food-seeking behavior, standard chow was removed from the cages two hours before and two hours after each session. Sessions alternated daily, starting with the palatable food session, followed by the empty cage session the next day, and so on (food-empty-food-empty-food-empty). The post-test was then performed, and the place preference index was calculated as in the cocaine CPP test.

**Fig 3.**
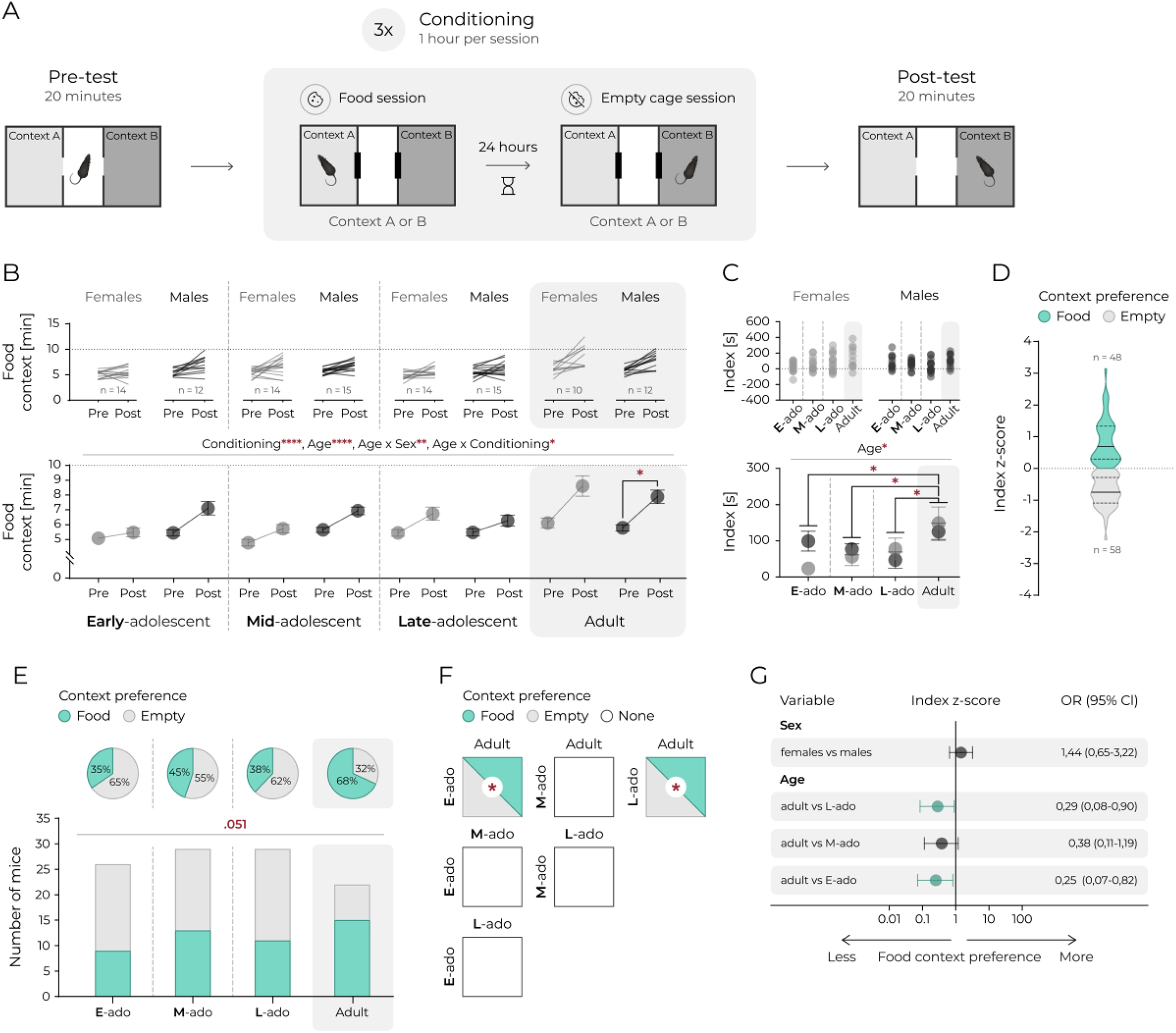
Preference for the context associated with palatable food is determined by age, not sex. **(A)** A schematic representation of the food CPP. **(B)** Time spent in the food context. Top panels: lines represent individual animals. Bottom panel: mean values. Circles and connecting lines represent means and matched values. Whiskers represent s.e.m. values. Dotted lines represent random value (i.e. 10 min). Females and males are shown in gray and black respectively. Statistical analysis was performed using 3-way ANOVA and post hoc Tukey’s HSD, ‘*’ corresponds to p ≤ .05, ‘**’, p ≤ .01, ‘****’ p ≤ .0001. **(C)** Difference in seconds between time spent in food context posttest and time spent in empty context pretest (index). Top panels: individual animals. Bottom panel: mean values. Whiskers represent s.e.m. values. Dotted lines represent no change. Statistical analysis was performed using 2-way ANOVA and post hoc Tukey’s HSD, ‘*’ corresponds to p ≤ .05. **(D)** Distribution of context preference (food or empty) in posttest. Sex and age data were combined. Index z-score calculated as: (index of individual animal – mean index of all mice) / standard deviation of index of all mice). Context preference interpreted according to index z-score value: positive, i.e. > 0 – “food” or negative, i.e. < 0 – “empty”. Solid line represents the median, dashed lines mark the quartiles. **(E)** Fractions of animals showing increase in preference of food context. Top panels (pie charts): percent of animals expressing food (green) or empty (white) context preference in each age group. Bottom panels: proportion of animals expressing food (green bars) or empty (white bars) context preference. χ^2^ test indicated a difference between proportions approaching statistical significance, p = .051. **(F)** Pairwise comparisons of the fraction of animals with increased food context preference at different development stages. Each square represents a comparison between each age group. The color of the square fill indicates the predominance of the proportion of animals expressing food (green), empty (gray) or none (white) preferences of one group over another. All pairwise χ^2^ comparisons were performed, ‘*’ corresponds to p ≤ .05. **(G)** Estimated effects of sex and age on the odds of increased preference for social context based on logistic regression. The circles and horizontal lines indicate the odds ratio and corresponding 95% confidence intervals. Statistically significant effects are marked green.

### Data analysis

All statistical analyses were performed using GraphPad Prism 10.4.2, except for four-way analysis of variance (ANOVA), which was conducted in R (4.5.0). Animals excluded from experiments based on pre-test criteria (those that spent more than 70% of time in any conditioning context) are listed in **Table S1**. No outliers were detected in the experimental data using the ROUT test. The results were analyzed with ANOVA followed by Tukey’s HSD. Additionally, χ2 was used for analyzing categorical data, and logistic regression assessed the effects of initial bias, sex, age, and reward type on preference or aversion of the conditioned context as a binary outcome. Correlations were evaluated using linear regression.

## Results

### Adolescent mice show decrease in social context preference

First, we examined the rewarding effects of social contact. We used the social CPP test (**Fig.1A**), where contact with a sibling of the same sex or social isolation is paired with two different types of novel bedding (social and isolation contexts, respectively). An increase in time spent in the social context from pre- to post-test is interpreted as evidence that an association has been learned between bedding and social contact. Initially, we analyzed the complete data set, excluding only animals with an initial preference for any context exceeding 70%. Mice spent significantly more time in the social context from pre- to post-test, and the analysis indicated a possible effect of age between males and females (**Fig.1B**, 3-way ANOVA: Fconditioning_(1.184)_ = 46.53, p < .0001, Fage_(3.184)_ = .67, p = .555, Fsex_(1.184)_ = 1.10, p = .295, Fage*sex_(3.184)_ = 3.13, p = .027; Fage*conditioning_(3.184)_ = 2.54, p = .058; Fsex*conditioning_(1.184)_ = .420, p = .518; Fage*sex*conditioning_(3.184)_ = 1.31, p = .273). To reduce the influence of initial context preference and simplify analysis, we used the social preference index, which measures the difference in time spent in the social conditioned context between post-test and pre-test (**Fig.1C**). We found that age significantly affected preference for the context associated with social reward (**Fig.1C**, 2-way ANOVA: Fage_(3.184)_ = 2.97, p = .033; Fsex_(1.184)_ = 0.34, p = .561; Fage*sex_(3.184)_ = 1.66, p = .179). There was no significant effect of sex nor interaction between sex and age in the analysis of variance. Although post-hoc comparisons did not reach statistical significance, we observed a trend toward reduced social CPP in late-adolescents compared to adults (**Fig.1C**), aligning partially with our recent results [40]. The effect of age was also evident when sex was excluded from the analysis to reduce complexity, and preference was classified as a binary outcome (i.e., z-score, see **Fig.1D**). This was assessed through cumulative frequency distributions of animals exhibiting a preference for the social context across different stages of adolescence (**Fig.1E**, χ^2^(3) = 11.44, p = .009). Mid- and late-adolescent mice showed significantly lower preference for the social context compared to adults (Fig.1E-F, 39% vs 71%, χ^2^(1) = 8.37, p = .004; 46% vs 71%, χ^2^(1) = 4.69, p = .030, respectively). Furthermore, the proportion of mid-adolescent mice that developed a preference for the social context was significantly lower than that of early-adolescent animals (**Fig.1E, F**, 39% vs 63%, χ^2^(1) = 5.70, p = .017). Correspondingly, when preference was evaluated as a binary outcome through logistic regression considering initial bias, sex, and age (**Fig. 1G**), there was a significant reduction in the odds of social context preference for mid-adolescent mice compared to adults (95% CI [0.08, 0.65], p = .007). A strong effect of initial preference on odds of expressing social CPP (odds ratio 95% CI [0.06, 0.23], p < .0001) was also observed.

As we [38] and others [34,36,47,48] have previously reported, CPP is confounded by reward-independent initial and acquired context preferences. While there was no initial context preference when analyzing the entire data set (cellulose preference 49. 95 ± 9%, one-sample t-test vs 50%, p > .05), there was a significant effect of sex on initial preference, as well as a significant interaction between sex and age (**Supplementary Fig.1A**, 2-way ANOVA Fage(3.21) = 1. 46, p = .230, Fsex_(1.21)_ = 10.82, p = .001, Fage*sex_(3.21)_ = 3.89, p = .009). The odds of cellulose bedding preference in the pretest was 29% higher for females compared to males (**Supplementary Fig.1B**, 95% CI [1.30, 4.10], p = .005) and 37% lower for late-adolescent mice compared to adults (95% CI [0.14, 0.97], p = .044). Additionally, as previously noted (**Fig. 1G**), the initial preference for the social context was the strongest predictor of conditioned preference. Indeed, the effects of conditioning were only significant in animals assigned the less preferred bedding as the social context (**Supplementary Fig.2A**, 4-way ANOVA significant effects: Finitialbias_(4.03)_ = 95.0, p < .001, Fconditioning_(3.36)_ = 52.7, p < .001, Fconditioning*initialbias_(3.36)_ = 59.4, p < .001, Fsex*age_(4.03)_ = 3.90, p < .01, Fsex*age*conditioning_(3.36)_ = 2.66, p < .01), despite no correlations between pre-test and post-test time in social context (r^2^ = 0.02, p > .05). Animals with a pre-test preference for the social context did not increase their preference from pre- to post-test, which is the primary criterion for successful conditioning-based learning. This reflects an inherent issue with the CPP task, and the benefits and equivalency of biased versus unbiased designs have been extensively discussed [36,47–49]. Therefore, to directly compare with the biased cocaine and palatable CPP tests, next we analyzed only the social CPP results from animals with no initial preference for the social context (i.e., a biased design).

There was a significant increase in time spent in the social context from pre- to post-test, with no effects of sex, age, or significant interactions (**Fig.1H**, 3-way ANOVA: Fconditioning_(1.105)_ = 111.6, p <.0001; Fage_(3.105)_ = .65, p = .585; Fsex_(1.105)_ = 2.10, p = .150; Fage*sex_(3.105)_ = .991, p = .400; Fage*conditioning_(3.105)_ = 1.14, p = .337; Fsex* conditioning_(1.105)_ = .072, p = .788; Fage*sex*conditioning_(3.105)_ = .273, p = .845). We found no significant effect of sex or age on social contact preference (**Fig.1I**, 2-way ANOVA: Fage_(3.104)_ = 1.58, p = .199; Fsex_(1.104)_ = .22, p = .638; Fage*sex_(3.104)_ = .28, p = .841). The proportion of animals that increased their preference for the social context was mostly similar to that in the unbiased analysis (**Fig.1J**), although fewer significant effects were observed. While cumulative frequency distributions of animals preferring the social context were not significantly different in adolescence (**Fig.1K**, χ² test, p > .05), a higher proportion of late-adolescent mice preferred the social context compared to mid-adolescent mice (63% vs. 37%, χ² (1) = 3. 96, p = .046). No significant effects were found in the logistic regression analysis, with one result nearing significance (**Fig.1M**, adult vs. mid-adolescent, p = .059). Overall, these findings mirror the trends observed in the unbiased analysis, although with fewer significant effects due to the smaller sample size. In summary, both unbiased and biased analyses highlight two separate effects: a lower preference for social-conditioned context in adolescent mice compared to adults, possibly varying between different stages of adolescence, and a qualitative difference in the proportion of animals that develop a preference for the social context between developmental stages.

### Age has no significant effect on cocaine conditioned place preference

Given these findings, we next investigated whether the reduction in social CPP observed during adolescence reflects a social-specific phenomenon or a broader attenuation of reward learning across modalities. We began with cocaine-induced CPP, as this paradigm is well-established and cocaine is a drug of abuse known to produce robust CPP with relatively few confounding or aversive effects. The rewarding effects of cocaine in adolescent and adult mice were assessed using a three-compartment CPP test (**Fig. 2A**). As expected, a significant increase in time spent in the context associated with cocaine injection was observed in female and male mice (**Fig.2B**, 3-way ANOVA: Fconditioning_(1.83)_ = 67.53, p < .0001, Fage_(3.83)_ = 6.26, p = .001, Fsex_(1.83)_ = .21, p = .648, Fage*sex_(3.83)_ = .309, p = .819, Fage*conditioning_(3.83)_ = .64, p = .590, Fsex*conditioning_(1.83)_ = .218, p = .642, Fage*sex*conditioning_(3.83)_ = 0.61, p = .614). There was a significant effect of age on the increase in preference, however, it was not observed when the preference index was assessed (**Fig.2C**, 2-way ANOVA: Fage_(3.83)_ = .64, p = .590, Fsex_(1.83)_ = .22, p = .642, Fage*sex_(3.83)_ = .61, p = .614). The absence of age-related differences in cocaine reward preference is further supported when preference is defined as a binary outcome (**Fig.2D**), indicating comparable proportions of animals exhibiting cocaine CPP across developmental stages (**Fig.2E**, χ^2^ test, p > .05), and no difference in pairwise comparisons (**Fig.2F**, all χ^2^ test, p > .05). Accordingly, logistic regression showed no significant effects of age or sex on preference (**Fig.2G**).

### Palatable food preference is lower in adolescent mice

Next, we assessed CPP using other natural reward, palatable food, with a similar setup as with cocaine CPP (**Fig.3A**). This method followed the procedure described by Clough and colleagues [46], but without an extended restriction of chow access. We found that palatable food significantly increased preference for the associated context, with a notable interaction between age and sex and between age and conditioning (**Fig.3B**, 3-way ANOVA: Fconditioning_(1.98)_ = 81.01, p < .0001, Fage_(3.98)_ = 12.08, p < .0001, Fsex_(1.98)_ = 3.52, p = .064, Fage* sex_(3.98)_ = 5.49, p = .002, Fage*conditioning_(3.98)_ = 3.65, p = .015, Fsex*cond_(1.98)_ = .34, p = .562, Fage*sex*cond_(3.98)_ = 1.82, p = .149). Preference for the food context, measured as the index, was significantly affected by age but not sex or age and sex interaction (**Fig.3C**, 2-way ANOVA: Fage_(3.98)_ = 3.65, p = .015, Fsex_(1.98)_ = .34, p = .642, Fage*sex_(3.88)_ = 1.82, p = .149). Early-, mid- and late-adolescent mice showed significantly lower conditioned preference than adults (**Fig.3C**). Preference categorized as a binary outcome (**Fig.3D**) revealed, that effect of age was also nearly statistically significant in the cumulative frequency analysis that excluded sex factor (**Fig.3E**, χ^2^(1) = 3.795, p = .051). Pairwise comparisons of the fractions that increased preference for the food-conditioned preference revealed significant differences between E-ado or L-ado mice and adult animals (**Fig.3E&F**, 35% vs 68%, χ^2^(1) = 8. 37, p = .004, 38% vs 68%, χ^2^(1) = 4. 69, p =.030). These effects remained significant when assessed with logistic regression (**Fig. 3G**). The odds of preferring the food context were lower by 25% for early-adolescent (95% CI [.07,.82], p = .026) and by 28% for late-adolescent mice (95% CI [.08,.90], p = .036) relative to adults. The time spent in the reward context during pre-test correlated with the time in post-test for both cocaine (r^2^ = .17, p < .0001) and palatable food CPP (r^2^ = .10, p = .001). Motor activity did not confound the results in any CPP task (see **Supplementary Fig.3A-D**).

### Adolescent mice show decreased reward place preference compared to adults

Lastly, we have performed a direct comparison of conditioned preference for all types of rewards, using normalized z-scores of the preference indices to exclude confounding effects of the differences in the post-test length and ignoring the effect of sex to simplify the analysis. There was a significant overall effect of age on conditioned preference; however, no effect of reward type or interaction between age and type of reward (**Fig.4A**, 2-way ANOVA: Freward_(2.30)_ = .064, p = .938, Fage_(3.30)_ = 3.21, p = .023, Freward*age_(6.30)_ = 1.07, p = .381). Adult mice had higher reward preferences than mid- and late-adolescent mice (**Fig.4A**). Accordingly, when we performed a logistic regression analysis for the effects of reward type, the age effect was robust, as the odds of preferring reward context was decreased by 48% for mid-adolescent (**Fig.4B**, 95% CI [0.24, 0.92], p = .028) relative to adult mice, given that the other variables in the model are held constant. Finally, we note that there was no overall effect of sex or reward type on the expression of the conditioned preference. In order to provide context for the observed changes, we present in **Fig.4C** the trend for reward CPP against puberty onset, the reward system development time-course, and vulnerability to mental disorders (based on Uhlhaas and collaborators [5]). While this is a speculative analysis, we note that the peak of developmental changes coincides with the observed decrease in CPP in mid-adolescence.

**Fig 4.**
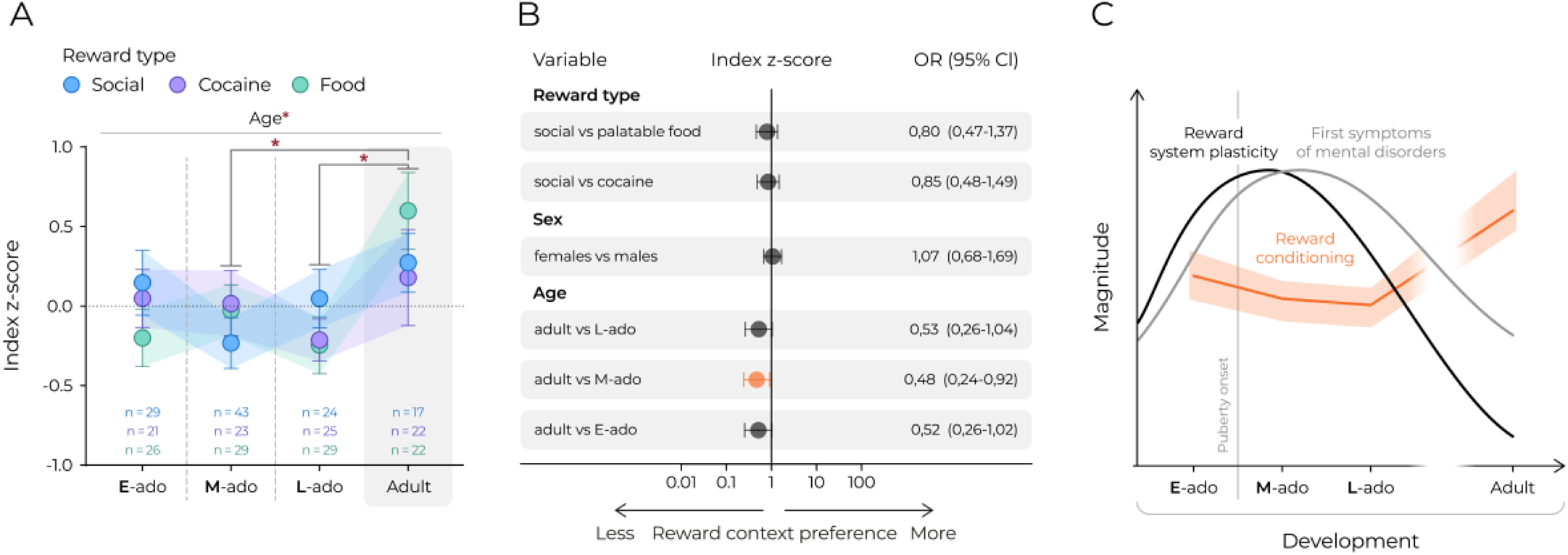
Adolescent mice express decreased reward CPP than adults, but greater plastic potential for reward system. **(A)** Comparison of relative conditioned context preference between social, cocaine and food reward. Sex data were combined. Index z-score calculated for each CPP experiment separately as: (index of individual animal – mean index of all mice) / standard deviation of index of all mice). The colored circles represent mean values, the color-corresponding shadows and whiskers represent s.e.m. values. Statistical analysis was performed using 2-way ANOVA and post hoc Tukey’s HSD, ‘*’ corresponds to p ≤ .05. **(B)** Estimated effects of reward type, sex and age on the odds of increased preference for reward context based on logistic regression. The circles and horizontal lines indicate the odds ratio and corresponding 95% confidence intervals. Statistically significant effects are marked orange. **(C)** A graphical representation of the developmental trajectories of social and non-social conditioned place preference in mice. The orange curve with the color-corresponding shadow represent mean and s.e.m. of reward conditioning (based on **Fig 4A**). The black and gray lines indicate the sensitive periods for reward system and emergence of first symptoms of mental disorders in humans, respectively (data based on [5]).

## Discussion

We find that the expression of reward-conditioned preference in adolescent mice is weaker compared to adults, and this effect was mostly consistent regardless of the type of stimulus or sex. This is an unexpected observation, as there is conflicting data for social CPP in mice [28], and the lower preference is intuitively at odds with the reported heightened reward sensitivity in adolescence [4,20]. Place preference is the most established method to evaluate the rewarding effects of a stimulus. Therefore, the reproducibility and comparability of CPP experiments have been extensively discussed, as even slight differences in protocol, such as experimental setup or behavioral parameters, can significantly impact results [34,47,48,50]. Notably, initial context bias plays a key role in forming conditioned place preference [36,48]. In theory, an unbiased design can detect both rewarding and aversive qualities of the stimulus. However, because rodents show varying CPP strength across different stimuli [51], pre-existing contextual biases may mask subtle behavioral changes. We previously used an unbiased design for the social CPP task [38,40,41], but employed biased designs for drug-conditioned CPP [44]. The original palatable-food CPP design is biased [46], and we adopted this design in the method used here. To prevent confounding bias effects in comparisons between rewards, we include both biased and unbiased analyses of social CPP and find that their results are consistent regardless of the approach. The main behavioral measure is the change in time spent in the conditioned context from pre-test to post-test, with z-score normalization applied to account for differences in test duration. Importantly, all testing was conducted under minimal stress conditions with mice bred on-site, and we assume their affective states were consistent across all experiments. We should also note that no monitoring of the estrous cycle was performed, which is a significant limitation for interpreting sex effects. The influence of female sex hormones on cocaine reward is well-known, with estrogen enhancing and progesterone reducing reward sensitivity [52,53]. Although juvenile females do not yet experience regular cycles, it remains possible that adult females could show increased cocaine CPP if conditioning occurs during hormonally sensitive cycle phases. Lastly, we are cautious in interpreting reward-type effects on CPP. As mentioned in the Results, the experiment was not designed to compare sensitivity to social and nonsocial rewards, and the differences in tasks may substantially influence conditioning. For cocaine, dose-response relationship was not investigated, so we cannot rule out the possibility that different doses could lead to a different increase in context preference. Similarly, in the case of palatable food, the period of limited chow access prior to testing would strongly influence motivation to seek food, and we deliberately limited deprivation. For these reasons, we avoid interpreting the effects of reward type and focus instead on sex and age effects that are not test-type dependent.

In line with our findings, Murray et al. reported a similar mid-adolescent decrease in social reward motivation in rats using an operant conditioning paradigm, although they observed the highest motivation in early adolescent animals [33]. Conversely, Nardou and colleagues [28] have elegantly demonstrated in the social CPP task that social contact is rewarding in male and female adolescent but not adult mice. We have previously shown that we could fully replicate these findings [38]. However, when we increased the number of conditioning sessions and ensured that social contact was with siblings, social CPP was strongly expressed by 14-week-old adult mice. This aligns with the role of social context, prior social experiences, internal state, and kinship in social CPP, which we and others have highlighted [38,40,42,54]. Additionally, we note that human adolescents show transient increases in social motivation toward non-kin peers rather than family members [4,20]. This may explain why social CPP could be enhanced during adolescence when conditioned with unfamiliar conspecifics. Similar to Murray et al. [33] and studies in mice [28,42], we observed no sex differences in social conditioned behaviors, contradicting some earlier reports showing higher social CPP reward in male rats compared to females [54,55] and higher social CPP in female mice compared to males [54], depending on social housing and prior social isolation. Furthermore, prior research indicates that adolescent rodents often display increased intake and motivation for palatable food [29–32], especially during mid-puberty [30], which contrasts with our findings. Methodological differences, such as species, reward type, or housing conditions, limit direct comparison. Age-related differences in food CPP might also reflect variations in reward consumption under food-restricted conditions. Shteyn & Petrovich show that adult rats consistently prefer palatable food regardless of satiety, while adolescents do so only when satiated [32]. Adult mice have also demonstrated CPP for palatable food mixes, including chow, but not for chow alone [46], suggesting high-fat and sugar components might be the primary rewards. In our study, animals were food-deprived only for two hours before conditioning, so the mild hunger might cause adults to prefer more rewarding components, forming stronger context associations. Adolescents, meanwhile, may balance their intake more evenly, resulting in weaker CPP. We did not assess food intake, which is a limitation. We found no consistent sex effect on food CPP; differences between early adolescent females and males were due to females spending more time in the neutral middle compartment, likely related to anxiety-like behaviors. The lack of sex effects aligns with some previous studies [56,57], but not the recent reports showing stronger CPP in females [32,59,60]. Lastly, the increased drug reward sensitivity in adolescents is often linked to stress effects [25] or reduced sensitivity to drug aversiveness [13,60], but in this study, stress was minimized, and cocaine was chosen as the drug of abuse with the fewest confounding or aversive effects. Overall, although our findings differ from some previous reports, we believe most earlier studies were not explicitly designed to assess reward sensitivity. Despite these limitations, our results provide strong evidence for reduced CPP in adolescent compared to adult mice.

Four primary factors influence the change in stimulus-induced context preference: (1) reinforcement, which reflects the intrinsic rewarding properties of the stimulus; (2) motivational state, encompassing the animal’s drive for reward-seeking; (3) memory, involving the acquisition and recall of the context-reward association; and (4) the conditioned response, representing behavioral changes following conditioning [61]. The individual contributions of these factors and their potential interactions cannot be fully disentangled in the CPP outcome. Given this complexity, the reduced CPP observed during adolescence could reflect several underlying mechanisms, including a diminished perceived value of the stimuli, impaired reward learning, or a shift toward increased novelty-seeking. First, it should be stressed that the lower increase in preference is unlikely to result from impaired learning. As previously discussed, our earlier studies demonstrated that juvenile mice develop social CPP following shorter conditioning protocols than those required for adults [38,40], suggesting that adolescents exhibit comparable or even enhanced learning efficiency. This interpretation is further supported by our current and prior findings on cocaine-induced CPP [39], which was consistently robust across all age groups, indicating intact reward-based learning in adolescents. Second, we found no evidence of a shift toward novelty-seeking behavior overriding conditioned responses in adolescent mice. In both the cocaine and food CPP paradigms, juvenile mice did not show a significant post-conditioning increase in time spent in the neutral context. Although they spent more time in the neutral zone overall compared to adults, this pattern was independent of conditioning and did not coincide with increased exploratory behavior, as the total distance traveled during post-tests consistently increased with age. Collectively, these findings support the interpretation that the attenuated CPP observed during adolescence is more likely due to decreased reward sensitivity or motivation rather than deficits in learning or an increase in novelty-seeking behavior.

From the basic perspective of Skinner’s behavioral learning theory [62,63], behavior is shaped through reinforcement and punishment. Therefore, we propose that the reward stimuli employed in our experiments may have been less reinforcing or the absence of reward less punishing for adolescent mice compared to adults, resulting in attenuated learning through conditioning. The developmental changes in reward system responsivity may underline this age-related difference in conditioned place preference. We observed a general trend of increased reward CPP from adolescence to adulthood, while the neural plasticity of the reward system follows an opposite pattern [4,5]. Mid-adolescence, characterized by peak developmental remodeling, may show the most consistent decrease in reward sensitivity across different reward types. In humans, adolescence involves a greater influence of emotional biases on executive functions [20,64]. Executive functions can be divided into “cold,” which involve cognitive control and logical reasoning, and “hot,” which are emotionally driven and linked to impulsivity and reward sensitivity, peaking in mid-adolescence [64]. It is plausible that under emotionally neutral conditions, reward-related stimuli are perceived as less salient, resulting in decreased reward sensitivity at certain adolescent stages, and thus a weaker expression of reward preference. This could serve as an adaptive mechanism, preventing unnecessary plastic changes in the reward system unless triggered by emotionally salient or socially relevant events. Such gating may protect against maladaptive changes that contribute to the development of disorders of affect and motivation, which often emerge during adolescence [18,29]. We hypothesize that adolescence is a developmental window of reduced reward sensitivity unless stimuli are emotionally or socially meaningful. Future research should investigate how social context, stress exposure, hormonal status, and individual behavioral profiles influence reward sensitivity and vulnerability to psychopathology during adolescence. Ultimately, these insights could guide the development of targeted pharmacological and psychotherapeutic treatments for mental illnesses that arise during adolescence and persist into adulthood.

## Acknowledgments

The social and cocaine CPP experiments were supported by grant OPUS 2019/35/B/NZ7/03477 and the palatable food experiment was supported by PRELUDIUM-21 2022/45/N/NZ4/01504, both from the National Science Centre Poland and the statutory funds of the Maj Institute of Pharmacology of the Polish Academy of Sciences in Krakow.

## Data availability statement

All data are available at Zenodo 10.5281/zenodo.16753864. Raw video recordings and Med-PC data will be made available at reasonable request.

---

**Table S1.** Age, weight and number of excluded animals in all CPP tests.

| CPP test | Sex | Age bin | Excluded n* | Tested n | Subject's weight at post-test [g] |  |  | Subject's age at post-test (days) |  |  |
| --- | --- | --- | --- | --- | --- | --- | --- | --- | --- | --- |
|  |  |  |  |  | Range | Mean | SEM | Range | Mean | SEM |
| social (unbiased) | females | Early-adolescent | 1 | 18 | 4,5 | 13,9 | 0,3 | 2 | 31 | 0,1 |
|  |  | Mid-adolescent | 0 | 32 | 4,5 | 15,2 | 0,2 | 5 | 37 | 0,4 |
|  |  | Late-adolescent | 0 | 20 | 2,7 | 15,5 | 0,2 | 4 | 44 | 0,4 |
|  |  | Adult | 1 | 16 | 4,1 | 19,8 | 0,3 | 2 | 82 | 0,2 |
|  | males | Early-adolescent | 1 | 20 | 4,2 | 17,6 | 0,3 | 2 | 32 | 0,1 |
|  |  | Mid-adolescent | 3 | 44 | 8,8 | 17,4 | 0,4 | 4 | 37 | 0,3 |
|  |  | Late-adolescent | 0 | 30 | 6,7 | 19,1 | 0,3 | 6 | 42 | 0,3 |
|  |  | Adult | 0 | 12 | 3,9 | 23,7 | 0,3 | 8 | 83 | 0,9 |
| social (biased) | females | Early-adolescent | - | 11 | 5,2 | 16,3 | 0,7 | 3 | 32 | 0,3 |
|  |  | Mid-adolescent | - | 19 | 7,4 | 17,1 | 0,6 | 5 | 37 | 0,5 |
|  |  | Late-adolescent | - | 17 | 7,2 | 18,3 | 0,6 | 6 | 42 | 0,4 |
|  |  | Adult | - | 9 | 2,9 | 20,1 | 0,3 | 2 | 82 | 0,2 |
|  | males | Early-adolescent | - | 18 | 7,4 | 16,1 | 0,5 | 2 | 32 | 0,1 |
|  |  | Mid-adolescent | - | 24 | 11,7 | 15,6 | 0,5 | 4 | 36 | 0,3 |
|  |  | Late-adolescent | - | 7 | 6,3 | 18,0 | 1,0 | 5 | 43 | 0,6 |
|  |  | Adult | - | 8 | 3,9 | 23,6 | 0,5 | 8 | 83 | 1,2 |
| cocaine | females | Early-adolescent | 1 | 12 | 4,5 | 11,8 | 0,3 | 2 | 31 | 0,2 |
|  |  | Mid-adolescent | 0 | 12 | 3,3 | 14,5 | 0,3 | 1 | 38 | 0,1 |
|  |  | Late-adolescent | 0 | 13 | 2,0 | 14,9 | 0,2 | 2 | 44 | 0,2 |
|  |  | Adult | 1 | 10 | 2,6 | 20,8 | 0,2 | 3 | 118 | 0,4 |
|  | males | Early-adolescent | 0 | 9 | 5,5 | 13,1 | 0,6 | 1 | 31 | 0,2 |
|  |  | Mid-adolescent | 0 | 11 | 2,6 | 17,9 | 0,3 | 3 | 38 | 0,4 |
|  |  | Late-adolescent | 0 | 12 | 5,4 | 18,8 | 0,5 | 1 | 43 | 0,1 |
|  |  | Adult | 2 | 12 | 7,5 | 25,5 | 0,6 | 6 | 112 | 0,7 |
| palatable food | females | Early-adolescent | 0 | 14 | 3,8 | 11,9 | 0,3 | 2 | 31 | 0,2 |
|  |  | Mid-adolescent | 1 | 14 | 2,5 | 14,6 | 0,2 | 0 | 38 | 0,0 |
|  |  | Late-adolescent | 0 | 14 | 4,1 | 16,3 | 0,3 | 2 | 44 | 0,2 |
|  |  | Adult | 2 | 10 | 4,6 | 20,7 | 0,4 | 2 | 115 | 0,3 |
|  | males | Early-adolescent | 2 | 12 | 7,7 | 13,6 | 0,7 | 2 | 31 | 0,2 |
|  |  | Mid-adolescent | 0 | 15 | 4,9 | 16,4 | 0,4 | 0 | 38 | 0,0 |
|  |  | Late-adolescent | 0 | 15 | 2,8 | 18,9 | 0,2 | 1 | 45 | 0,1 |
|  |  | Adult | 0 | 12 | 6,2 | 28,1 | 0,6 | 1 | 102 | 0,1 |

\*Number of mice that did not meet criterion of spending less than 70% of time in any of conditioning contexts

**Table S2.**
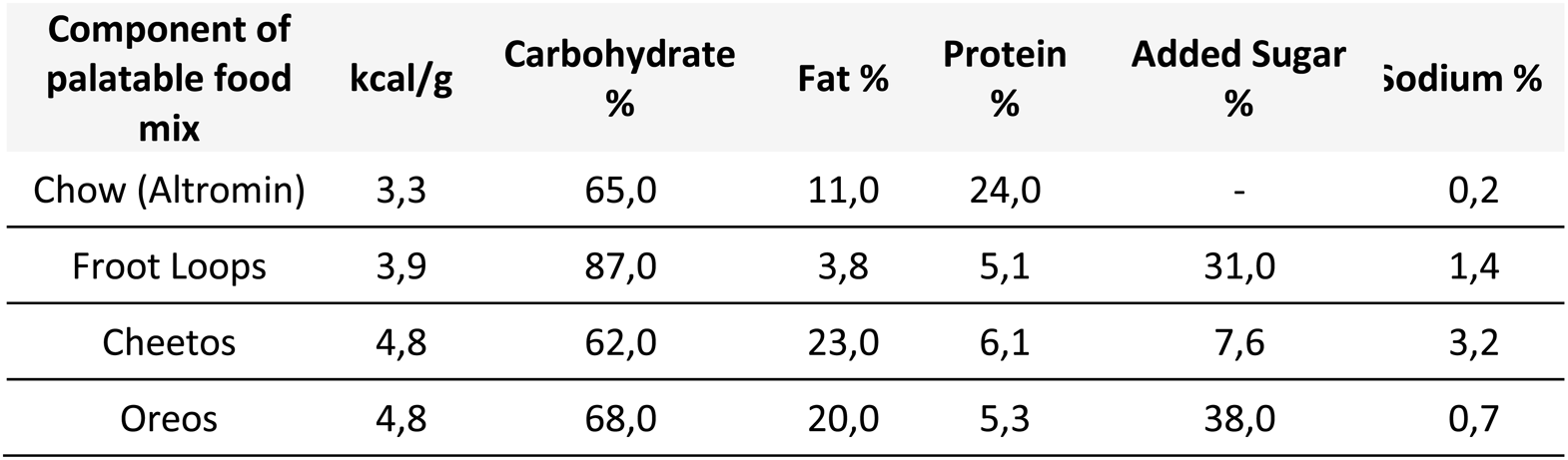
Nutritional comparison of ingredients of palatable food mix used in food CPP test.

**Supplementary Figure.1.**
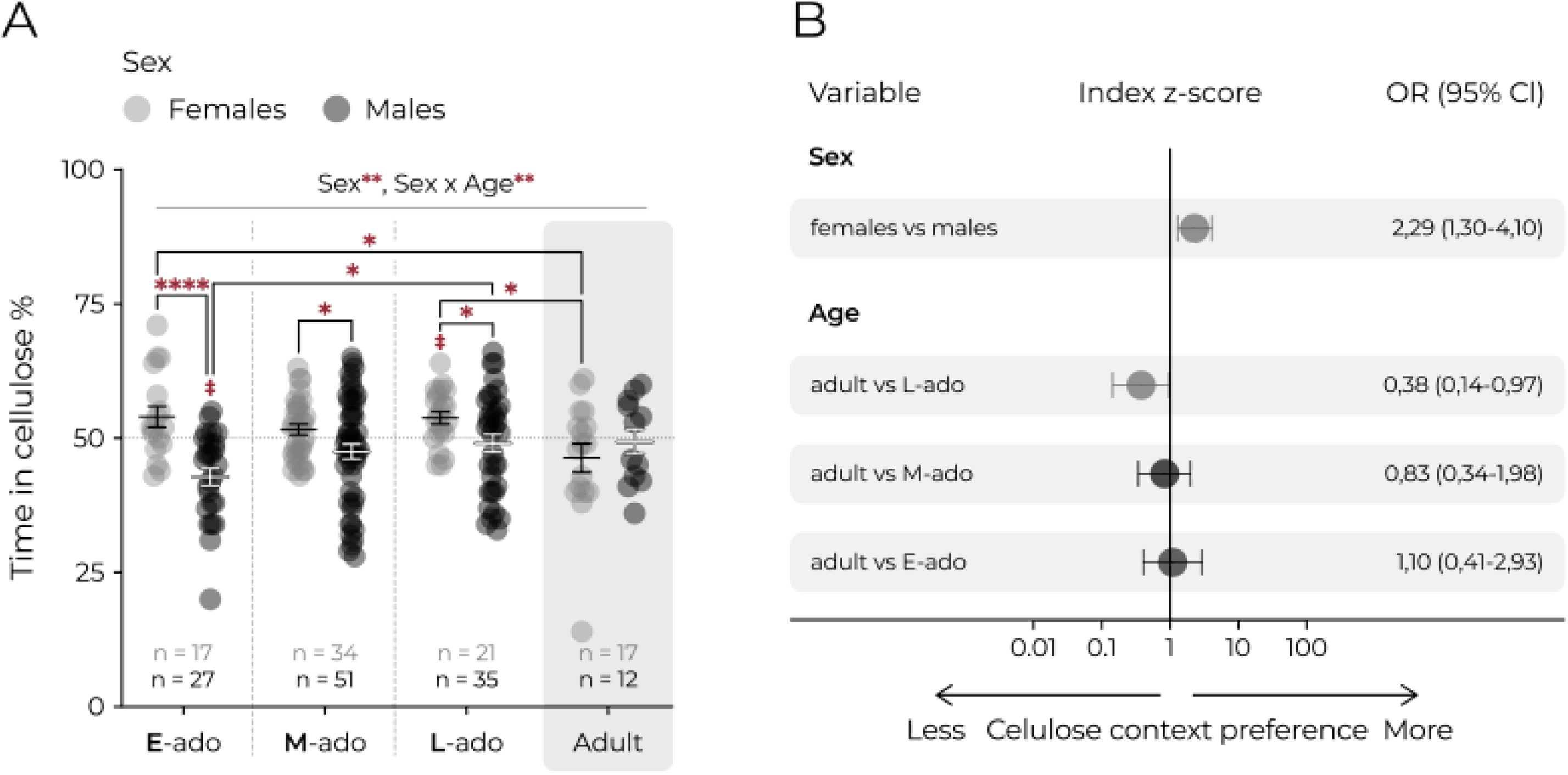
Mice express significant bias in initial preference in pretest before place conditioning. **(A)** Time spent in the cellulose context in pretest. Graph includes data from all animals, including animals later excluded based on criterion of initial context bias (>70% of time in any of the contexts in pretest). Each circle represents an individual animal. Whiskers represent mean and s.e.m. values. Dotted line represents random value (i.e. 50%). Females and males are shown in gray and black respectively. Statistical analysis was performed with 2-way ANOVA: Fage_(3.21)_ = 1.46, p = .230, Fsex_(1.21)_ = 10.82, p = .001, Fage*sex_(3.21)_ = 3.89, p = .009 and Tukey’s HSD, “*” corresponds to *p ≤ .05, **p ≤ .01, and ****p ≤ .0001 and “^#^”, p ≤ .08. **(B)** Estimated effects of sex and age on the odds of increased preference for cellulose context based on logistic regression. The circles and horizontal lines indicate the odds ratio and corresponding 95% confidence intervals. Statistically significant effects are marked gray.

**Supplementary Figure.2.**
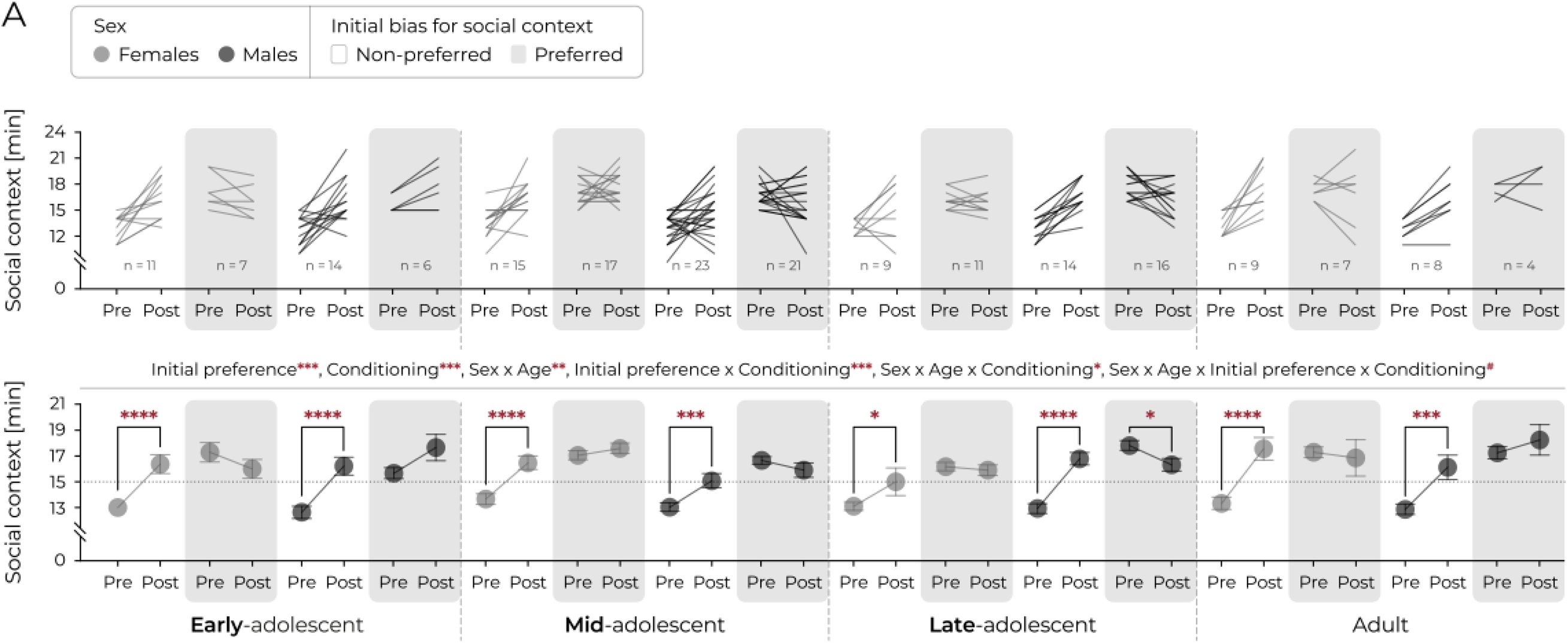
Increase in place preference after conditioning differs as a function of initial bias for social context. **(A)** Time spent in the social context, according to initial bias. Top panels: lines represent individual animals. Bottom panel: mean values. Circles and connecting lines represent means and matched values. Whiskers represent s.e.m. values. Dotted lines represent random value (i.e. 15 min). Females and males are shown in gray and black respectively. Females and males are shown in dark gray and black respectively. Bias for social context for initially non-preferred and preferred context are marked white and light gray background, respectively. Statistical analysis was performed using 4-way ANOVA with matched values: : Finitialbias_(4.03)_ = 95.0, p < .001, Fconditioning_(3.36)_ = 52.7, p < .001, Fsex_(4.03)_ = .49, p = .486, Fage_(4.03)_ = 1.02, p = .386, Fsex*age_(4.03)_ = 3.90, p = .010, Fsex*initialbias_(4.03)_ = 1.47, p = .227, Fage*initialbias_(4.03)_ = .04, p = .990, Fsex*conditioning_(3.36)_ = .41, p = .523, Fage*conditioning_(3.36)_ = 1.85, p = .140, Fconditioning*initialbias_(3.36)_ = 59.44, p < .001, Fsex*age*initialbias_(4.03)_ = .58, p = .630, Fsex*age*conditioning_(3.36)_ = 2.66, p = .050, Fsex*initialbias*conditioning_(3.36)_ = .06, p = .806, Fage*initialbias*conditioning_(3.36)_ = .63, p = .599, Fsex*age*initialbias*conditioning_(3.36)_ = 2.61, p = .053, and post hoc Tukey’s HSD, ‘*’ corresponds to p ≤ .05, ‘**’, p ≤ .01, ‘***’, p ≤ .001, ‘****’ p ≤ .0001.

**Supplementary Figure.3.**
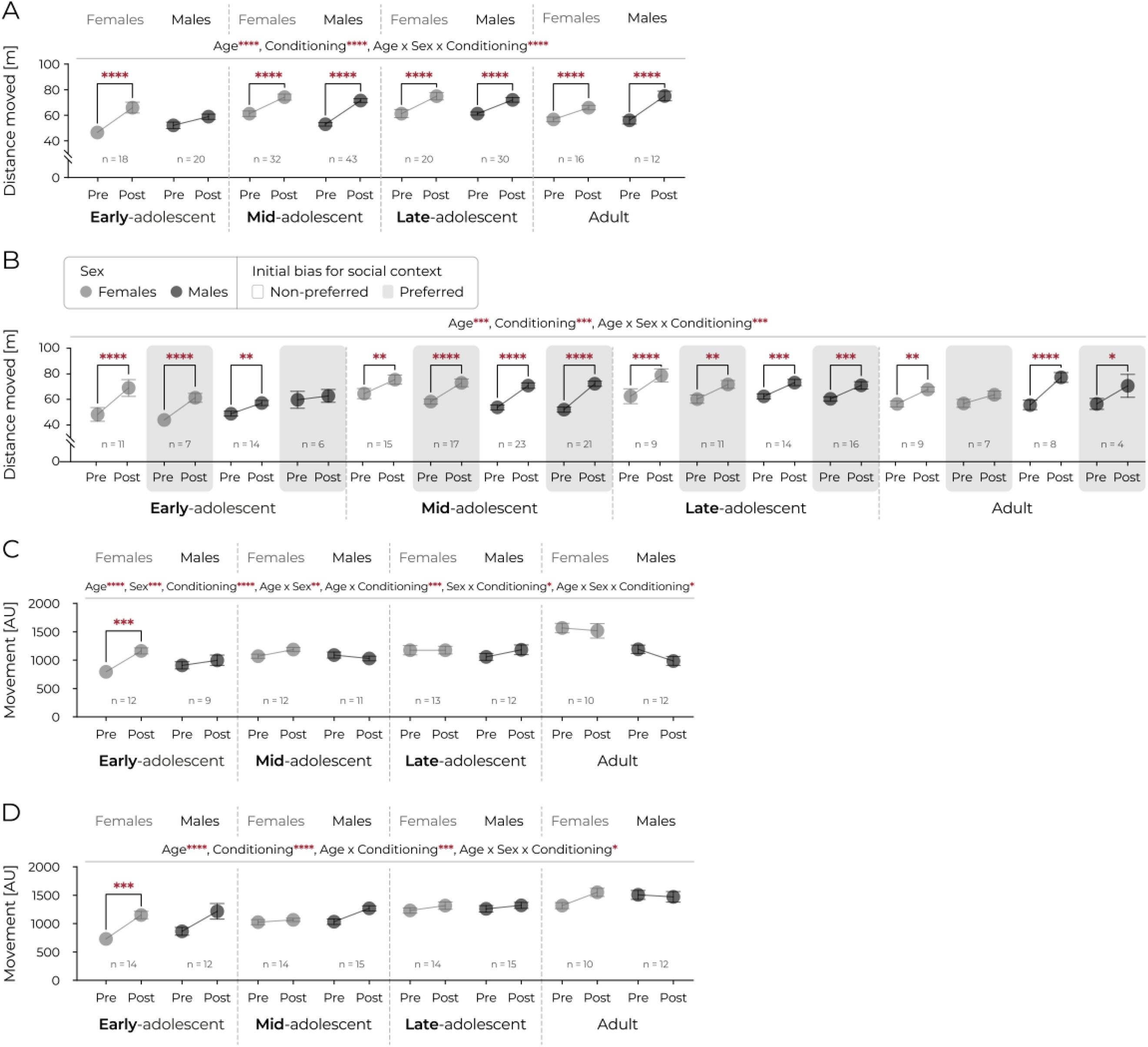
Analysis of motor activity during the pre-test and posttest phases in social, cocaine, and food CPP paradigms revealed no evidence of a confounding effect on context preference. **(A)** Distance moved during the pre- and post-test session in unbiased social CPP. Circles and connecting lines represent means and matched values. Whiskers represent s.e.m. values. Females and males are shown in gray and black respectively. Statistical analysis was performed using 3-way ANOVA with matched values was performed: Fconditioning_(1.18)_ = 242.4, p < .0001, Fage_(3.18)_ = 11.16, p < .0001, Fsex_(1.18)_ = .35, p = .554, Fage*sex_(3.18)_ = 1.80, p = .147, Fage*conditioning_(3.18)_ = 1.11, p = .346, Fsex*conditioning_(1.18)_ = .420, p = .994, Fage*sex*conditioning_(3.18)_ = 7.45, p < .0001 and post hoc Tukey’s HSD, ‘****’ corresponds to p ≤ .0001. **(B)** Distance moved during the pre- and post-test session, according to initial bias in social CPP. Circles and connecting lines represent means and matched values. Whiskers represent s.e.m. values. Females and males are shown in dark gray and black respectively. Bias for social context for initially non-preferred and preferred context are marked white and light gray background, respectively. Statistical analysis was performed using 4-way ANOVA with matched values: Fconditioning_65.64)_ = 214.03, p < .0001, Fage_(184.40)_ = 9.62, p < .001, Fsex_(184.40)_ = .09, p = .759, Finitialbias_(184.40)_ = 1.26, p = .263, Fsex*age_(184.40)_ = 2.06, p = .111, Fsex*initialbias_(184.40)_ = 2.63, p = .107, Fsex*conditioning_(65.64)_ = .06, p = .812, Fage*conditioning_(65.64)_ = 1.17, p = .321, Fconditioning*initialbias_(65.64)_ = 1.58, p = .210, Fage*sex*conditioning_(65.64)_ = 7.06, p < .001, Fsex*initialbias*conditioning_(65.64)_ = .00, p < .953, Fage*initialbias*conditioning_(65.64)_ = 1.63, p < .183, Fage*sex*initialbias_(184.40)_ =.94, p = .420, Fage*sex*initialbias*conditioning_(65.64)_ = .22, p = .885 and post hoc Tukey’s HSD, ‘*’ corresponds to p ≤ .05, ‘**’, p ≤ .01, ‘***’, p ≤ .001, ‘****’ p ≤ .0001. **(C)** Distance moved during the pre- and post-test session in cocaine CPP. Circles and connecting lines represent means and matched values. Whiskers represent s.e.m. values. Females and males are shown in gray and black respectively. Statistical analysis was performed using 3-way ANOVA with matched values was performed: Fconditioning_(1.18)_ = 3.62, p = .060, Fage_(3.83)_ = 10.78, p < .0001, Fsex_(1.18)_ = 12.33, p = .001, Fage*sex_(3.83)_ = 5.43, p = .002, Fage*conditioning_(3.82)_ = 7.69, p = .0001, Fsex*conditioning_(1.18)_ = 5.82, p = .018, Fage*sex*conditioning_(3.83)_ = 3.03, p = .034 and post hoc Tukey’s HSD, ‘*’ corresponds to p ≤ .05, ‘**’, p ≤ .01, ‘***’, p ≤ .001, ‘****’ p ≤ .0001. **(D)** Distance moved during the pre- and post-test session in food CPP. Circles and connecting lines represent means and matched values. Whiskers represent s.e.m. values. Females and males are shown in gray and black respectively. Statistical analysis was performed using 3-way ANOVA with matched values was performed: Fconditioning_(1.98)_ = 39.48, p < .0001, Fage_(3.98)_ = 27.29, p < .0001, Fsex_(1.98)_ = 3.20, p = .076, Fage*sex_(3.98)_ = .33, p = .801, Fage*conditioning_(3.98)_ = 6.77, p = .0003, Fsex*conditioning_(1.98)_ = .59, p = .445, Fage*sex*conditioning_(3.98)_ = 2.83, p = .042 and post hoc Tukey’s HSD, ‘*’ corresponds to p ≤ .05, ‘***’, p ≤ .001, ‘****’ p ≤ .0001.

**Supplementary Figure.4.**
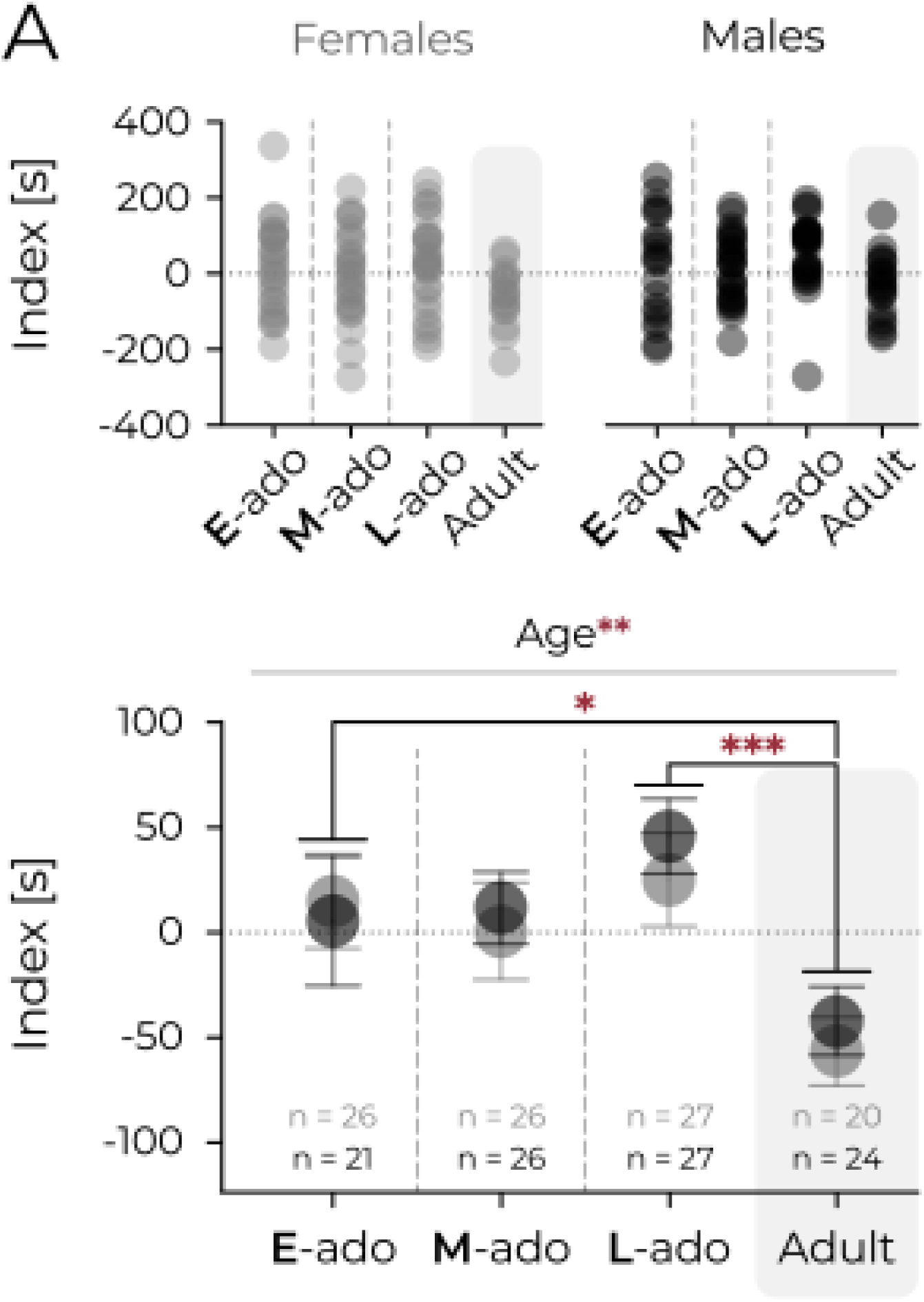
Adult mice, regardless of sex, spent less time in the neutral context than juvenile mice after conditioning. **(A)** Difference in seconds between time spent in neutral context posttest and time spent in neutral context pre-test (index). Cocaine and food CPP data were combined. Top panels: individual animals. Bottom panel: mean values. Whiskers represent s.e.m. values. Dotted lines represent no change. Statistical analysis was performed using 2-way ANOVA: Fage_(3.19)_ = 5.43, p = .0001, Fsex_(1.19)_ = .37, p = .542, Fage*sex_(3.19)_ = .18, p = .907 and post hoc Tukey’s HSD, ‘*’ corresponds to p ≤ .05, ‘**’, p ≤ .01.

